# LinearRaccess: linear-time approximation of RNA accessibility without base-pair span limits

**DOI:** 10.64898/2026.11.04.764167

**Authors:** Takumi Otagaki, Kengo Sato

**Affiliations:** School of Life Science and Technology, Institute of Science Tokyo, Room 401, Midorigaoka Building 6, Ookayama Campus, 2-12-1-M6-12 Ookayama, Meguro-ku, Tokyo 152–8550, Japan

**Keywords:** RNA secondary structure, accessibility, beam pruning, inside–outside algorithm, linear-time algorithm

## Abstract

**Motivation:** RNA accessibility is the probability that every nucleotide in a contiguous interval is simultaneously unpaired; it is widely used to predict binding sites for RNA-binding proteins and other RNA–RNA interactions, often in the form of the opening energy, an equivalent measure on a logarithmic scale. Existing tools restrict the maximum base-pair span, discarding long-range base pairs that matter biologically. Others sample structures, introducing statistical variance, or compute accessibility exactly without a span limit, at cubic time cost. No existing method computes the accessibility of windows longer than one nucleotide deterministically in linear time while preserving long-range base pairs. We present LinearRaccess, which computes accessibility directly in linear time via beam-pruned inside–outside dynamic programming with a difference-array aggregation step, without restricting base-pair span.

**Results:** For a sequence of length *N* and a fixed beam width *b*, LinearRaccess runs in *O*(*N* ) time and *O*(*Nb*) memory. Against exact accessibility on 12,872 bpRNA-1m(90) sequences of 100–1,000 nt, its median opening-energy error is below 0.001 kcal/mol (0.14 kcal/mol for 700–1,000 nt), and it distinguishes windows of very small accessibility, including 35-nt windows, below the resolution of sampling-based estimates. LinearRaccess analyzes the full 29,903-nt SARS-CoV-2 genome in 419 seconds.

**Conclusion:** LinearRaccess is the first method to compute the accessibility of windows longer than one nucleotide deterministically (without sampling) in linear time without restricting base-pair span.

**Availability:** https://github.com/TakumiOtagaki/LinearRaccess, archived at Zenodo (https://doi.org/10.5281/zenodo.23598502).

## 1. Introduction

Computational prediction of RNA secondary structure from nucleotide sequences is a long-standing problem in bioinformatics, with applications ranging from non-coding RNA annotation to therapeutic target identification [4], [15], [19], [20]. The classical approach based on thermodynamic models, particularly the nearest-neighbor model [15], formulates structure prediction, for a sequence of length *N*, as a free energy minimization problem solved by dynamic programming in O(*N* ^3^) time and O(*N* ^2^) memory [24]. Since a single minimum-free-energy structure cannot capture the full ensemble of thermodynamically plausible structures, partition-function-based methods instead compute the Boltzmann-weighted ensemble of all possible structures, yielding base-pairing probabilities [16]. Base-pairing probabilities are widely used for assessing structural confidence and for other downstream structural analyses.

Beyond base-pair probabilities, RNA structure analysis also characterizes *accessibility* : the joint probability that all nucleotides in a contiguous interval [*s, e*] are simultaneously unpaired in the ensemble. Accessibility helps to predict binding sites for RNA–RNA and RNA–protein interactions: antisense oligonucleotides [4], miRNAs [9], and RNA-binding proteins [11] tend to bind accessible regions. RNA–RNA interaction predictors such as RNAup [17] and IntaRNA [13] incorporate target-site accessibility directly into their interaction energy models as an *opening energy*, the free energy required to keep the site unpaired (Eq. 2 below). Candidate sites are then compared and ranked by the sum of this opening energy and the hybridization energy. Existing tools for computing accessibility include RNAplfold [1], [2] and Raccess [9], which compute the accessibility of a local window, often imposing a maximum base-pair span *W* to achieve O(*NW* ^2^) time complexity. While this approach is computationally tractable, it discards long-range base pairs, which can play a biological role. Theoretical and experimental studies indicate that RNA secondary structure can bring the 5^*1*^ and 3^*1*^ regions of viral genomes and mRNAs into close spatial proximity [10], [21], via base pairs that are inherently long-range. Computing accessibility with the inside–outside algorithm without a span limit avoids this restriction, but has been limited to cubic time and quadratic memory [2].

Inspired by LinearFold [7], which computes minimum free energy structures in linear time via beam pruning, several methods have extended this paradigm to ensemble-based quantities. LinearPartition [23] approximates the partition function and base-pairing probabilities in linear time while preserving long-range base pairs. CapR [5] computes per-nucleotide structural context profiles (the probability of being in each of six structural contexts: paired, plus the five unpaired contexts below), revealing binding specificities of RNA-binding proteins; LinearCapR [18] extends this computation to linear time. LinearSampling [22] performs stochastic structure sampling in linear time, and can estimate accessibility by counting unpaired windows in sampled structures, though this sampling-based approach introduces statistical noise, especially for rare events. These methods share the key insight that beam pruning—keeping only the *b* most promising states at each position—dramatically reduces computational cost while preserving accuracy. A separate line of work, DeepRaccess [6], predicts accessibility for a single window length (35 nt for the published model) directly with a neural network trained on Raccess output.

However, a notable gap remains: while base-pair probabilities and structural context profiles can be computed deterministically in linear time, no method computes the accessibility of windows longer than one nucleotide deterministically in linear time while preserving long-range base pairs.

Here we present LinearRaccess, which fills this gap by computing accessibility deterministically in linear time using beam-pruned inside–outside dynamic programming combined with a difference-array aggregation, without imposing the global base-pair span limit that local folding approaches require.

We validate LinearRaccess against the Raccess reference, characterize its accuracy against window-limited Raccess and RNAplfold on 12,872 bpRNA sequences in both probability and opening energy (Table 1), compare it with LinearSampling and DeepRaccess, and measure its runtime and memory scaling on the bpRNA dataset, together with a whole-genome runtime measurement on the SARS-CoV-2 genome.

**Table 1.** Accuracy of LinearRaccess (beam=200), Raccess Window (*W* =100), and RNAplfold (*W* =100) against Raccess Exact on *n* bpRNA sequences per half-open length bin [*·, ·*) (last bin closed), over all windows of length *w* ∈ *{*1, 3, 7, 10, 35*}*. All methods use Turner 1999 parameters. Absolute error is |*P*_approximate_ *− P*_exact_| for accessibility probabilities, and opening-energy error is 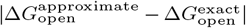 in kcal/mol (Eq. 2); each median pools all (position, window-length) cells across the sequences in that bin.

|  |  | Median absolute probability error |  |  | Median opening-energy error (kcal/mol) |  |  |
|---|---|---|---|---|---|---|---|
| Length range (nt) | *n* | LinearRaccess | Raccess Window | RNAplfold | LinearRaccess | Raccess Window | RNAplfold |
| 100–300 | 11,102 | 2.2×10<sup>−9</sup> | 5.6×10<sup>−5</sup> | 5.0×10<sup>−3</sup> | <0.001 | 0.18 | 0.87 |
| 300–500 | 1,442 | 6.9×10<sup>−7</sup> | 6.6×10<sup>−4</sup> | 1.1×10<sup>−2</sup> | 0.003 | 0.74 | 1.41 |
| 500–700 | 190 | 6.3×10<sup>−6</sup> | 7.6×10<sup>−4</sup> | 1.7×10<sup>−2</sup> | 0.024 | 0.72 | 1.56 |
| 700–1,000 | 138 | 1.4×10<sup>−4</sup> | 1.7×10<sup>−3</sup> | 1.8×10<sup>−2</sup> | 0.136 | 0.67 | 1.23 |
| **All (100–1,000)** | **12,872** | **1.0×10<sup>−8</sup>** | **1.5×10<sup>−4</sup>** | **7.2×10<sup>−3</sup>** | **<0.001** | **0.34** | **1.04** |

## 2. Methods

### 2.1 Problem definition

Given an RNA sequence *x* = *x*_0_*x*_1_ … *x*_*N−*1_ of length *N*, we consider the Boltzmann ensemble of pseudoknot-free secondary structures. Accessibility of interval [*s, e*] (inclusive) is defined as:

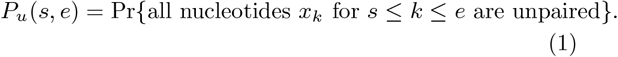

Figure 1 illustrates this quantity for a target window [*i, j*]. In general, 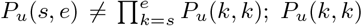 *P*_*u*_(*k, k*) is the pernucleotide unpaired probability, the special case of Eq. 1 for a single position. For a fixed window length *w* = *e − s* + 1, LinearRaccess computes *P*_*u*_(*i, i*+*w−*1) for every valid starting position *i*. Accessibility is often reported instead as the opening energy

**Fig. 1.**
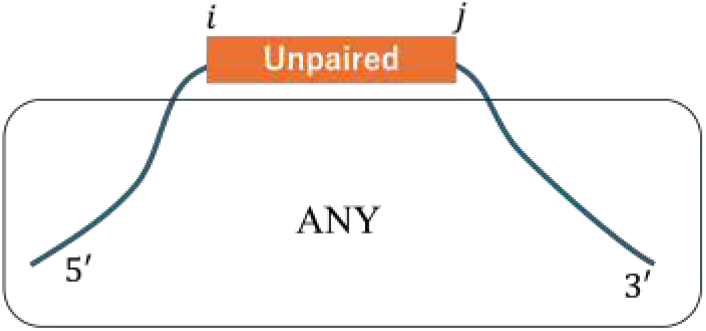
Accessibility is the probability that every nucleotide in a target window [*i, j*] is unpaired, with no constraint on the rest of the structure.

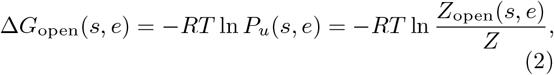

where *R* is the gas constant, *T* the absolute temperature, *Z* the partition function, and *Z*_open_(*s, e*) the partition function restricted to structures leaving [*s, e*] unpaired; this is Raccess’s default output [9], and LinearRaccess reports either quantity. Since Δ*G*_open_ is logarithmic in *P*_*u*_, windows with very small accessibility (e.g. *P*_*u*_ *≈* 10^*−*6^ versus 10^*−*8^) remain comparable on this scale, although their differences in *P*_*u*_ are negligible. We therefore evaluate accuracy on both scales (§3).

### 2.2 Beam-pruned inside–outside algorithm

LinearRaccess builds on the beam-pruned inside–outside framework shared with LinearPartition and LinearCapR. The algorithm maintains a set of states representing partial parse trees of the RNA structure – equivalent to parsing under a context-free grammar for RNA secondary structure [7], with each retained state (*t, i, j*) pairing a non-terminal type *t* (e.g., a base pair closing *i* and *j*) with a span [*i, j*], and an associated partition function *Q*_*t*_(*i, j*) representing the total Boltzmann weight of all structures on the subsequence *x*_*i*_*x*_*i*+1_ … *x*_*j*_ under state *t*; beam pruning bounds the number of retained states at each position, keeping the total work linear in *N*.

The key insight is that, for any single nucleotide position *k*, the probability distribution over the *maximal unpaired segment* containing *k* – a run of consecutive unpaired positions [*s, e*] such that *x*_*s−*1_ and *x*_*e*+1_ (if they exist) are each paired, not necessarily to each other – can be computed from the inside partition functions *Q*_*t*_(*i, j*) together with outside partition functions that score how a span is enclosed by the rest of the structure. Let *C* = *{*exterior, hairpin, bulge, internal, multiloop*}* denote the five *unpaired* structural contexts such a segment can occur in, defined by the loop that encloses it in the nearest-neighbor model: the exterior loop (not enclosed by any base pair), a hairpin loop (closed by a single base pair with no pair inside), a bulge or an internal loop (between a closing base pair and a single inner base pair, unpaired on one side or on both sides), or a multiloop (closed by one base pair and containing two or more others). The sixth, paired context (a base pair itself) cannot contain an unpaired segment, so it plays no role here. For each context *c* ∈ *C*, define

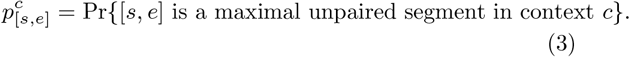

The algorithm has two stages. The first is a beam-pruned inside–outside pass (Algorithm 1) that scores partial structures and computes the segment probabilities 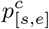 defined above, following the parsing framework shared with LinearFold, LinearPartition, and LinearCapR. The second is a difference-array aggregation (Algorithm 2) that turns these segment probabilities into window accessibilities in linear time, using the difference-array technique that LinearCapR [18] applies to per-nucleotide aggregation. This retargeting, from per-nucleotide marginals to joint window probabilities, is this paper’s main technical contribution.

#### Algorithm 1

Beam-pruned inside pass (schematic)

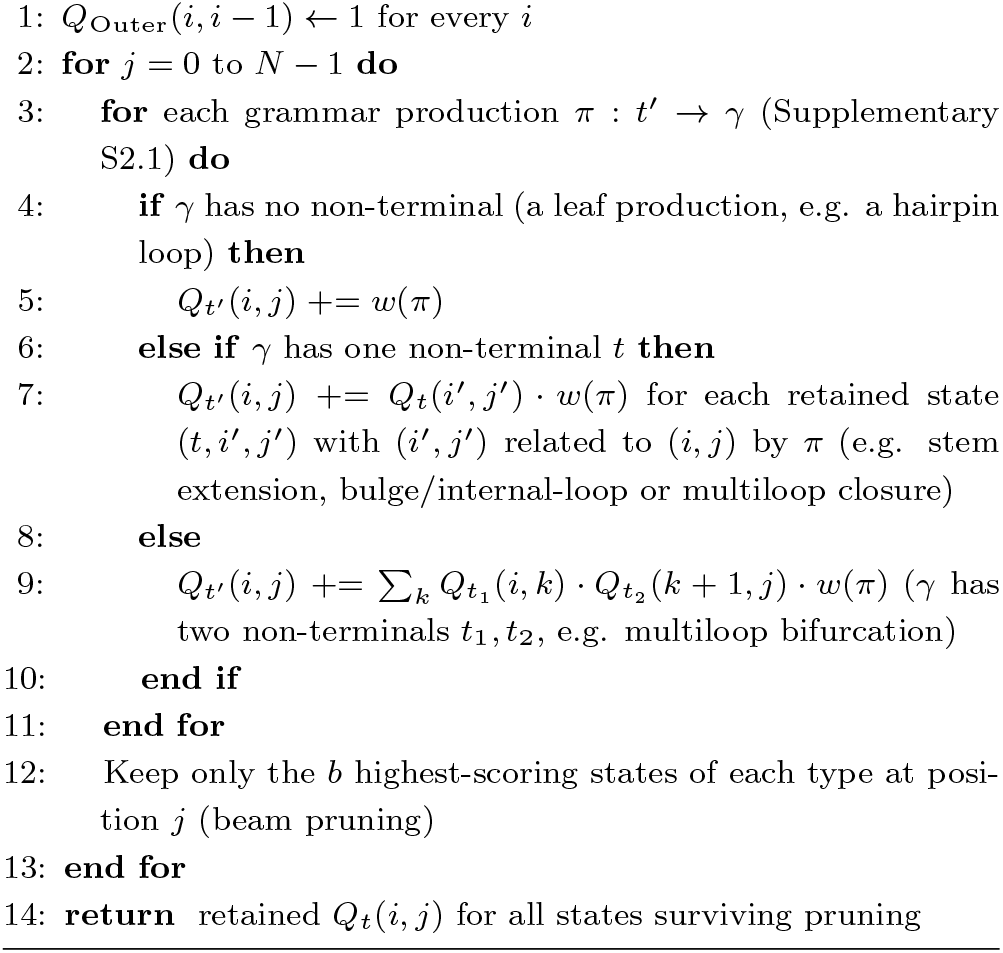

The outside pass traverses the same retained states in reverse to score how each span is enclosed by the rest of the structure, and the five-context decomposition combines the inside and outside scores into the per-context segment probabilities 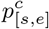 above. This inside–outside computation and five-context decomposition are not novel to this paper: LinearCapR [18] already introduces the same recurrences, as a per-nucleotide marginal; LinearRaccess reuses them, computing them per segment [*s, e*] instead. Algorithm 1 is deliberately abstract; *t, t’*^*1*^ range over the full seven-state machine (Outer, Stem, StemEnd, Multi, MultiBif, Multi1, Multi2), *π* ranges over that grammar’s productions, and 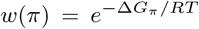 is *π*’s Boltzmann weight; Supplementary Information S2 gives the complete production list, citing LinearCapR for the production weights and the complete inside and outside recurrences.

Every unpaired nucleotide belongs to exactly one maximal unpaired segment, and that segment occurs in exactly one context *c* ∈ *C*; a window [*i, i*+*w −*1] is entirely unpaired exactly when it falls inside one such segment. Summing over every such occurrence gives

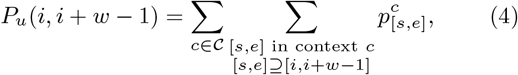

which is exhaustive and has no double counting: in any single structure in which the window is unpaired, it lies in exactly one maximal unpaired segment, of exactly one context, so the events summed are disjoint. Write

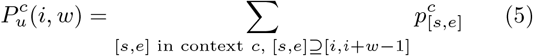

for the inner sum above, the accessibility of the window in context *c*, so that 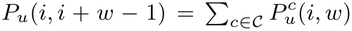; this window-level quantity, not 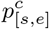 itself, is what Algorithm 2 below actually computes.

Each such segment [*s, e*] is relevant to a window [*i, i*+*w−*1] only if the window lies entirely inside it, i.e. [*s, e*] ⊇ [*i, i*+*w −*1]; we add its probability to a difference array once, over the range of window-start positions it covers, and recover all requested window accessibilities with a single prefix sum. This applies to the four loop-interior contexts (hairpin, bulge, internal, multiloop); the exterior context, bounded by the sequence ends rather than by enclosing structure, is instead computed directly from the boundary partition functions at each window start.

#### Algorithm 2

Difference-array window aggregation

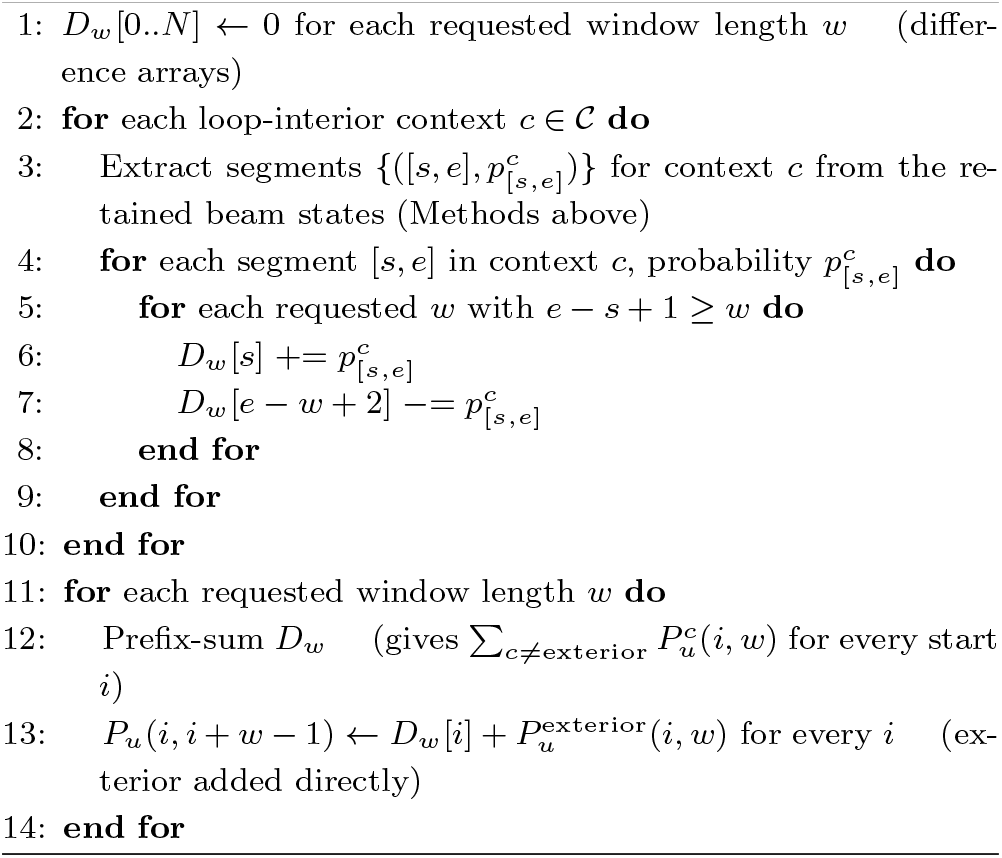

The difference-array aggregation (Algorithm 2) turns an aggregation that would otherwise cost O(*N* ) per event into O(1) per event plus one O(*N* ) prefix sum per window length, independent of how many windows the event covers.

The beam-pruned inside–outside pass costs 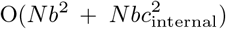 . The lengths of unpaired runs in hairpin loops and in multiloops are bounded by loop caps *c*_hairpin_ and *c*_multi_. The O(*Nb*^2^) term covers multiloop bifurcations that combine two retained states, and multiloop unpaired runs of length up to *c*_multi_ at each of the *b* retained states; hairpin loops add only O(*Nc*_hairpin_). The 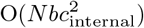 term accounts for internal/bulge loop closures, whose two flanking unpaired runs are jointly bounded by the standard ViennaRNA [12]/Turner `MAXLOOP` convention, *c*_internal_ = 30 nt. By default the hairpin and multiloop caps follow the beam, *c*_multi_ = *c*_hairpin_ = *b*. The only fixed structural limit is then this conventional internal-loop bound, and *b → ∞* recovers the uncapped model; setting beam=0 disables pruning altogether (exact mode). Algorithm 2 extracts each of the 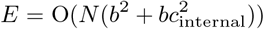 segments 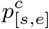 once and adds it, in O(1) each, to the difference arrays of the *K* requested window lengths it can contain; one O(*N* ) prefix sum per window length then recovers all window accessibilities. The overall time complexity is therefore 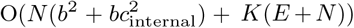, linear in *N* for any fixed beam width *b* (default 200) and number of window lengths *K*.

## 3. Results

We evaluated LinearRaccess on four aspects: exact-mode validation against Raccess reference, approximation accuracy, comparison with the sampling-based LinearSampling and the neural-network DeepRaccess, and long-sequence scaling (including a SARS-CoV-2 whole-genome runtime measurement). Unless stated otherwise, LinearRaccess runs with its defaults (beam *b* = 200, loop caps *c*_multi_ = *c*_hairpin_ = *b*), and every experiment requests the window lengths *w* ∈ *{*1, 3, 7, 10, 35*}*; *w* = 35 is the window length of DeepRaccess’s published model and exceeds the 30-nt internal-loop bound. Throughout, we compare against two reference configurations: *Raccess Exact*, run without a span limit as ground truth, and *Raccess Window* (*W* =100), run with Raccess’s default maximum base-pair span of *W* = 100 nt. For large-scale validation, we used 26,063 sequences from the bpRNA-1m(90) dataset [3], filtered to length *≥*50 nt and an ACGU-only alphabet, ranging from 50 to 4,381 nt. Comparisons with Raccess (§3.1, §3.2, the DeepRaccess comparison, and the SARS-CoV-2 beam-convergence check) use the Turner 1999 model, the energy model that both Raccess and LinearRaccess support; the LinearSampling comparison and the runtime measurements use LinearRaccess’s default Turner 2004 model. Unless noted otherwise, all runtime and memory measurements were single-threaded, on a single core of an AMD EPYC 9654 processor (TSUBAME4.0; GCC 11.5.0), for every tool compared.

### 3.1 Exact-mode validation

We compared LinearRaccess (beam=0, i.e., no pruning; loop caps = *∞*) against the Raccess reference implementation [9] (Supplementary S0) using the Turner 1999 energy model [14], on 300 randomly generated 100 nt RNA sequences, across window lengths *w* ∈ *{*1, 3, 7, 10, 35*}*. Across all 134,700 window-position combinations, the median probability error was 1.8*×*10^*−*8^, the 95th-percentile error was 5.2*×*10^*−*7^, and the maximum error was 1.6*×*10^*−*6^; the largest opening-energy difference was 5*×*10^*−*5^ kcal/mol, the precision of Raccess’s printed output. The Spearman rank correlation was 1 to within 10^*−*10^, confirming that Linear-Raccess correctly reproduces the reference ensemble when approximations are disabled.

### 3.2 Approximation accuracy

Practical use requires the approximations introduced in Methods (§2.2): beam pruning, with the hairpin and multi-loop caps following the beam. The comparison below uses the Turner 1999 energy model, matching Raccess. We additionally compare against RNAplfold [1], [2] (Supplementary S0), which estimates accessibility via local sliding-window folding averaged over every window containing the region, a different ensemble definition from Raccess Window’s single global ensemble. We restrict this comparison to the 12,872 sequences of 100–1,000 nt among §3.4’s dataset: shorter sequences trivially satisfy *N −* 1 *< W*, making Raccess Window identical to Raccess Exact, and Raccess Exact’s cubic cost limits it to 1,000 nt.

Across 12,872 bpRNA sequences ranging from 100 to 1,000 nt (Table 1), LinearRaccess is more accurate than Raccess Window in every length bin, on both scales. The pooled medians are dominated by sequences under 300 nt, on which beam pruning is rarely active (and on sequences no longer than the beam width, *N ≤* 200, LinearRaccess prunes nothing and is exact); the margin narrows for longer sequences (700–1,000 nt: 1.4*×*10^*−*4^ vs. 1.7*×*10^*−*3^ absolute, 0.14 vs. 0.67 kcal/mol), since more states compete for the fixed beam as sequences grow longer. The two longest bins have fewer sequences than the two shortest (190 and 138, versus 11,102 and 1,442), and their medians are therefore less statistically stable.

RNAplfold’s median error is larger than Raccess Window’s, despite both restricting base-pair span to *W* =100 and using Turner 1999 parameters, because RNAplfold averages local folding ensembles rather than computing a single global ensemble. Raccess Window’s error reflects its base-pair span limit, which LinearRaccess does not impose.

Median error understates a minority failure case for LinearRaccess, however: for 152 of the 12,872 sequences (1.2%), at least one position/window has an absolute error exceeding 0.5 (Figure 2 visualizes this spread against Raccess Exact); at beam=100 this rises to 789 sequences (6.1%). Supplementary Information S1 extends the error-distribution analysis, and S3 shows the error decreasing monotonically as the beam widens.

**Fig. 2.**
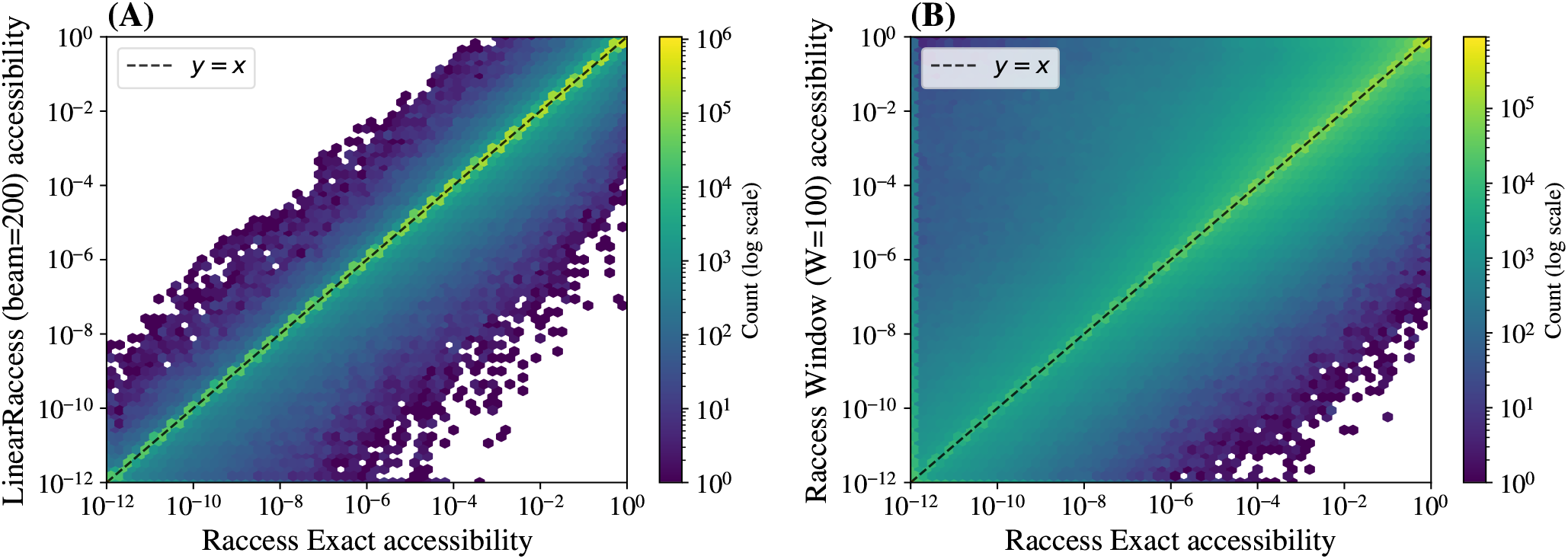
Per-cell accessibility values (12,872 bpRNA sequences, 100–1,000 nt, *w* ∈ {1, 3, 7, 10, 35} ) plotted against Raccess Exact, for LinearRaccess (beam=200; **A**) and Raccess Window (*W* =100; **B**). Color shows the count of cells per hexagonal bin (log scale); the dashed line marks exact agreement (*y* = *x*). LinearRaccess concentrates more tightly around the diagonal than Raccess Window, with a minority of off-diagonal cells corresponding to the beam-pruning failure case.

The absolute-error medians above are dominated by high-probability cells and are uninformative about relative accuracy where the true value is small, which is where the opening energy is informative. Restricted to the 1,489,567 cells where Raccess Exact’s accessibility falls between 10^*−*8^ and 10^*−*5^ (opening energies of 7–11 kcal/mol), LinearRaccess’s median relative error is below 0.01%, with a 90th percentile of 16%, and it returns a positive accessibility for all but 100 of the 11.4 million cells, all of them 35-nt windows with exact accessibility below 2 *×* 10^*−*20^.

### 3.3 Comparison with LinearSampling and Deep-Raccess

We compared LinearRaccess with LinearSampling [22], which estimates accessibility by counting unpaired windows in *k* sampled structures, and with DeepRaccess [6] (Supplementary S5 and S6). For LinearSampling we used *k* = 10^6^ samples, twenty times the 5 *×* 10^4^ used in its original accessibility evaluation, and the same Turner 2004 model as LinearRaccess; both were scored against LinearRaccess in exact mode (beam=0), which agrees with Raccess Exact (§3.1) and with LinearSampling’s own exact sampling (Supplementary S5). A window that is unpaired in none of the *k* samples is estimated as zero, and all such windows receive the same, infinite opening energy: the resolution of a sampling estimate is 1*/k*. On 20 sequences of 1,000–3,541 nt at beam width 500, 21% of the 10-nt windows and 95% of the 35-nt windows fell below it. This leaves LinearSampling’s within-sequence ranking of 35-nt windows almost uninformative (median Spearman correlation 0.16, versus 0.96 for LinearRaccess), whereas LinearRaccess returned a finite opening energy for every window of these sequences; Supplementary S5 compares the two methods’ accuracy across probability ranges, and their runtime.

The published DeepRaccess model predicts only *w* = 35 and was trained on the output of Raccess Window (*W* = 100) run with CONTRAfold parameters instead of Turner energies, a model that LinearRaccess does not support; we therefore scored each method against the model it approximates, on the 12,872 sequences of Table 1. Deep-Raccess’s median opening-energy error against its teacher was 0.53 kcal/mol (median within-sequence Spearman correlation 0.88), whereas LinearRaccess’s against Raccess Exact was below 0.001 kcal/mol at *w* = 35 (Spearman 1.00). LinearRaccess’s error grows with sequence length, however, and in the 700–1,000 nt bin exceeds DeepRaccess’s (0.62 vs. 0.39 kcal/mol). Because the two methods use different underlying energy models, these error values are not directly comparable as measures of physical accessibility accuracy. DeepRaccess was far faster: about 26-fold on one CPU thread, and all 12,872 sequences took 31 s on one GPU. DeepRaccess’s predictions also depend only on local subsequences: its teacher is itself an approximation that restricts base pairs to a span of *W* = 100, and for a sequence longer than 440 nt, DeepRaccess takes overlapping 440-nt subsequences as input, predicts each separately, and joins the results; it therefore represents neither long-range base pairs nor structure spanning more than one subsequence. LinearRaccess processes a whole sequence without either restriction, and handles any window length and either Turner model directly, whereas other window lengths or energy models require retraining DeepRaccess [6].

### 3.4 Long-sequence scaling

This section uses LinearRaccess’s default Turner 2004 model. Raccess does not support Turner 2004; Raccess Exact and Raccess Window below therefore run under Turner 1999.

Figure 3A shows the runtime scaling comparison across three methods on the bpRNA dataset; we measured the time for the same five requested window lengths used throughout this paper (*w* ∈ *{*1, 3, 7, 10, 35*}*). LinearRaccess (beam=200) ran on all 26,063 sequences and exhibits linear scaling (0.0115 seconds/nt). Raccess Exact shows cubic scaling; its O(*N* ^3^) cost limits it to sequences *≤*1,000 nt here. Raccess Window also scales linearly, with a smaller constant factor (0.0015 seconds/nt).

**Fig. 3.**
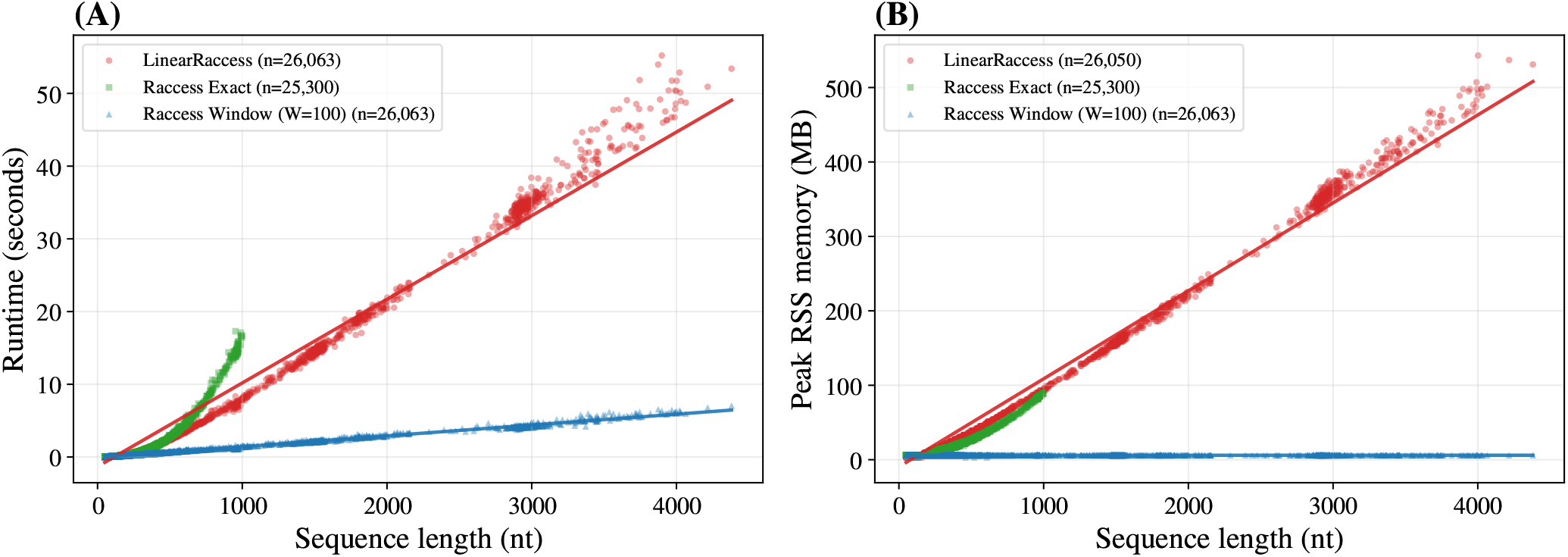
Runtime (**A**) and memory (**B**) on the bpRNA dataset [3] (26,063 sequences). LinearRaccess achieves linear scaling in both time and space, unlike Raccess Exact (run only up to 1,000 nt), which shows cubic time and quadratic memory growth. Raccess Window also scales linearly, but only because it restricts the maximum base-pair span; LinearRaccess achieves the same linear scaling without that restriction.

Figure 3B shows the corresponding memory scaling: LinearRaccess achieves linear memory scaling (0.12 MB/nt), requiring 531 MB for the longest sequence (4,381 nt). In contrast, Raccess Exact exhibits quadratic memory growth, consuming 89 MB for 1,000-nt sequences. Raccess Window achieves low, roughly constant memory usage (about 6 MB): its own *O*(*N* + *W* ^2^) complexity [9] is dominated by the fixed *W* ^2^ = 100^2^ term at these sequence lengths and barely grows with *N* . The linear memory scaling of LinearRaccess enables whole-genome analysis within the constraints of standard workstations (16 GB RAM). Raccess Exact’s quadratic scaling, projected to 30,000 nt, would need on the order of 80 GB.

### 3.5 Whole-genome runtime measurement: SARS-CoV-2

As a single further data point beyond the bpRNA range, LinearRaccess analyzes the full 29,903-nt SARS-CoV-2 genome (NC 045512.2) in 419 seconds and a measured peak memory of 4,404 MB under default parameters, some-what above what the linear fits on the shorter bpRNA sequences predict (344 seconds and 3.5 GB). To assess sensitivity to beam width at this length, we computed the genome’s ensemble free energy *G*_ensemble_, under the Turner 1999 model used for the accuracy comparisons above, across beam widths *b* ∈ *{*5, 10, 25, 50, 100, 200, 300, 500*}* (Figure 4). *G*_ensemble_ decreases across the tested beam widths, with changes of *−*1,381 kcal/mol from *b*=5 to 10, *−*182 kcal/mol from *b*=100 to 200, and *−*61 kcal/mol from *b*=300 to 500. The changes shrink as the beam widens, but *G*_ensemble_ has not converged by *b*=500, which already took 67 minutes and 10 GB; as for the longer bpRNA sequences (§3.2; Supplementary S3), a genome-length input leaves beam-pruning error that the default beam does not remove.

**Fig. 4.**
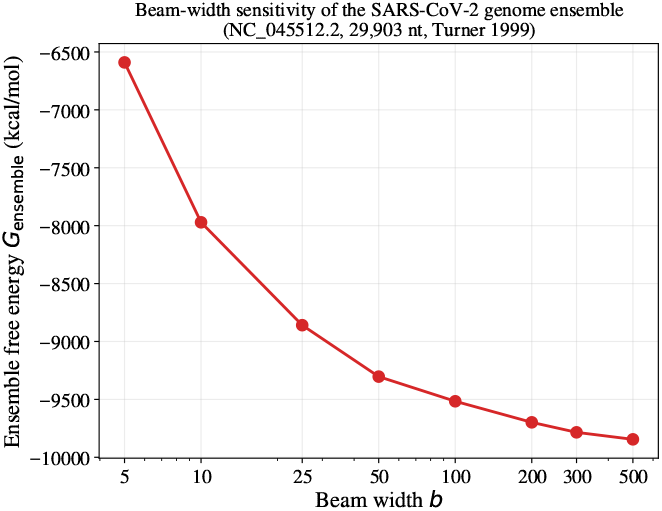
Ensemble free energy *G*_ensemble_ of the SARS-CoV-2 genome (NC 045512.2, 29,903 nt, Turner 1999 parameters) as a function of beam width *b*. Values decrease across the tested beam widths, with smaller changes at larger beams (text).

## 4. Discussion

For a user analyzing a full viral genome, LinearRaccess offers three practical advantages. Extrapolating Raccess Exact’s measured runtimes to whole-genome length gives one to five days (by the fitted power law, exponent 2.6, or by cubic growth), whereas LinearRaccess analyzes the full SARS-CoV-2 genome in 419 seconds (§3.5). It has no hard base-pair span limit, unlike Raccess Window, and processes a whole sequence at once, unlike DeepRaccess, which predicts long sequences from 440-nt subsequences (§3.3). As a deterministic computation, it also returns a finite opening energy for essentially every window, including windows too rarely accessible for sampling to resolve (§3.3), so that such windows can still be compared and ranked. These windows are not negligible: a site’s interaction free energy is the sum of its opening and hybridization energies, and a long binder such as an antisense oligonucleotide can offset opening energies well above sampling’s resolution, *RT* ln *k ≈* 8.5 kcal/mol at *k* = 10^6^; this matters most for long windows.

LinearRaccess also reports each window’s accessibility by loop context (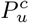, Eq. 5) at no extra cost, a decomposition RNAup also offers [17]. RNA-binding proteins differ in the contexts they prefer: at the same ACUK motif, an FMR1 isoform favors internal loops and bulges whereas FXR2 favors exterior and multiloop contexts [5], and HuR prefers exterior regions [8]. Since these context probabilities are sensitive to a base-pair span limit [5], and a sampling estimate resolves each context only from the samples in that context (Supplementary S7), deterministic, span-free loop-resolved accessibility may be useful for such analyses.

Accuracy under default settings is limited by beam pruning, and the required beam grows with both sequence length and window length (Supplementary S3, S4); for sequences beyond 1,000 nt and genome-length inputs, the beam should be widened where accuracy matters more than time.

## 5. Conclusion

LinearRaccess computes RNA accessibility in linear time without imposing a base-pair span limit, by combining beam-pruned inside–outside and difference-array aggregation algorithms (Algorithms 1–2). It may therefore be useful for accessibility-based analyses of long RNAs, such as RNA-binding-protein binding-site prediction on long non-coding RNAs, where long-range base pairs are common.

## Data Availability

LinearRaccess source code is available at https://github.com/TakumiOtagaki/LinearRaccess; the release used for every LinearRaccess result in this paper (v0.2.0) is archived at Zenodo, https://doi.org/10.5281/zenodo.23598502. Benchmark scripts, provenance manifests, and result bundles supporting the figures and tables in this paper are available at https://github.com/TakumiOtagaki/LinearRaccess-computational-experiments. Supplementary Information is available online.

## Acknowledgments

This study was carried out using the TSUBAME4.0 supercomputer at Institute of Science Tokyo.

