## Supplementary Information for "LinearRaccess: linear-time approximation of RNA accessibility without base-pair span limits"

### S0. Software versions and settings

All LinearRaccess results in this paper were generated with LinearRaccess v0.2.0 (git commit `0155fa4`, which differs from the tagged release `772f802` only in its version number; Zenodo DOI 10.5281/zenodo.23598502), with its defaults (beam 200, $c_{\text{multi}} = c_{\text{hairpin}} = b$) and window lengths $w \in \{1, 3, 7, 10, 35\}$. The SARS-CoV-2 ensemble free energies (main text §3.5) were computed with the preceding commit `0f78f13`; the later changes affect only the default beam width (set explicitly there) and the accessibility aggregation, not the ensemble free energy. The Raccess reference implementation (Kiryu et al., 2011) used throughout this paper is git commit `ad29d09`, which has no versioned release. RNAplfold (main text §3.2) is from ViennaRNA Package 2.7.2 (Lorenz et al., 2011), run as `RNAplfold -W 100 -L 100 -u 35 -P rna_turner1999.par`, explicitly selecting the bundled Turner 1999 parameter file so its energy model matches Raccess. LinearSampling (Zhang et al., 2023) (main text §3.3, S5) is git commit `1492a93`, run as `linearsampling -k 1000000 -b` $b$ with its built-in Turner 2004 parameters; window accessibility is the fraction of the $k$ sampled structures in which the window is unpaired. DeepRaccess (Hara et al., 2023) (S6) is git commit `1a37235` with the published `FCN_structured.pth` checkpoint (SHA-256 `136be4db...`), run with PyTorch 2.0.0 through its own preprocessing; its teacher, Raccess with CONTRAfold parameters, is the `run_raccess_contrafold` program of the same Raccess commit.

### S1. Beam-pruning failure case in the default-accuracy comparison

The main text (Results, §3.2) reports median errors and summarizes the beam-pruning failure case: 152 of the 12,872 sequences (1.2%) have at least one window with absolute error above 0.5 at the default beam=200, and 789 (6.1%) at beam=100. Figure S1 shows the full per-window error distribution underlying those medians, computed from all 12,872 bpRNA sequences (100–1,000 nt), split by sequence-length bin. Figure S2 shows, for the 789 sequences affected at beam=100, each sequence's maximum error at beam=100 and at beam=200; the median drops from 0.788 to 0.031.

Table 1 pools all five window lengths. Table S1 breaks its opening-energy error down by window length. For both methods and in every length bin, the error grows with $w$, since longer windows have smaller accessibilities, decided by rarer structures; LinearRaccess stays below Raccess Window at every $w$ and in every bin, but its margin narrows for the longest windows of the longest sequences (700–1,000 nt, $w = 35$: 0.62 vs. 1.41 kcal/mol).

Table S1: Median opening-energy error (kcal/mol) against Raccess Exact by window length, using Table 1's same 12,872 bpRNA sequences, length bins, and $n$ per bin.

| | LinearRaccess (beam=200) | | | | | Raccess Window ($W$=100) | | | | |
|---|---|---|---|---|---|---|---|---|---|---|
| Length range (nt) | $w$=1 | 3 | 7 | 10 | 35 | $w$=1 | 3 | 7 | 10 | 35 |
| 100–300 | <0.001 | <0.001 | <0.001 | <0.001 | <0.001 | 0.036 | 0.11 | 0.23 | 0.29 | 0.68 |
| 300–500 | <0.001 | 0.001 | 0.003 | 0.005 | 0.019 | 0.27 | 0.52 | 0.81 | 0.94 | 1.50 |
| 500–700 | 0.003 | 0.010 | 0.026 | 0.039 | 0.19 | 0.24 | 0.49 | 0.79 | 0.92 | 1.56 |
| 700–1,000 | 0.030 | 0.070 | 0.15 | 0.20 | 0.62 | 0.24 | 0.46 | 0.73 | 0.85 | 1.41 |
| **All (100–1,000)** | **<0.001** | **<0.001** | **<0.001** | **<0.001** | **<0.001** | **0.082** | **0.22** | **0.40** | **0.50** | **0.98** |

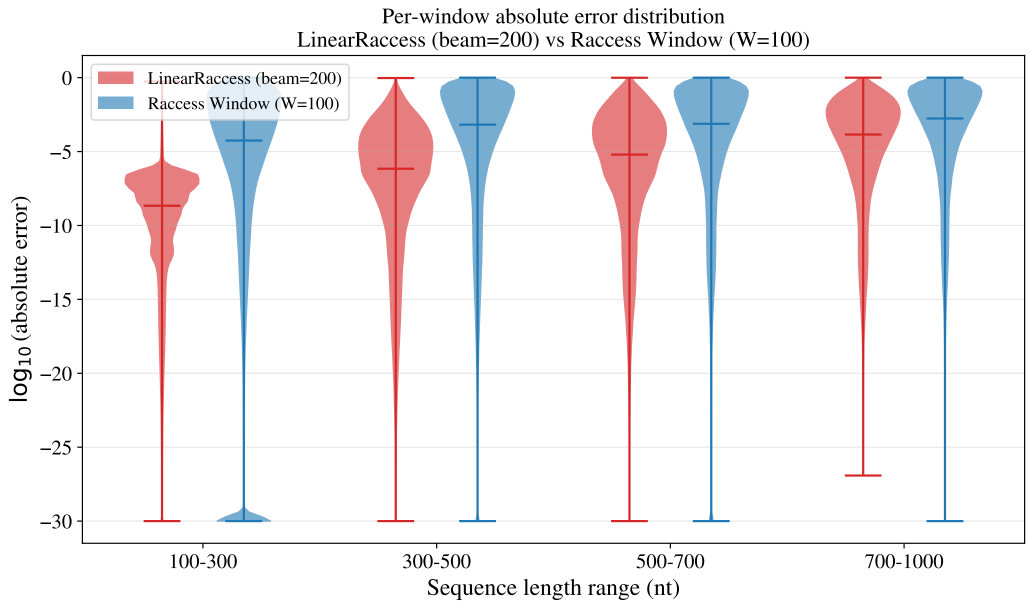


Figure S1: Per-window absolute-error distribution ($\log_{10}$ scale) of LinearRaccess (beam=200) and Raccess Window ($W$=100) against the exact reference, by sequence-length bin, over all windows of length $w \in \{1, 3, 7, 10, 35\}$. LinearRaccess's mass concentrates at much lower error than Raccess Window's; the failure case discussed in the main text is the minority of LinearRaccess windows in the thin tail near error $\approx 1$.

### S2. Full inside–outside recurrence and difference-array aggregation

Main-text Algorithm 1 is a schematic condensation: it writes every production abstractly as a leaf, one-non-terminal, or two-non-terminal rule over typed states $Q_t(i, j)$, and omits the outside pass entirely. This section gives the full seven-state grammar (S2.1), cites LinearCapR (Otagaki et al., 2026) for the inside–outside recurrence over that grammar rather than reproducing it (S2.2), and gives the segment-event extraction algorithm that turns those scores into $p^c_{[s,e]}$ (S2.3), this paper's own contribution. The grammar and non-terminal states (S2.1) are unchanged from LinearCapR. The segment-event extraction equations (S2.3) are adapted from LinearCapR's own published equations, re-labeled for this paper's $p^c_{[s,e]}$ notation and corrected in two places where LinearCapR's own per-nucleotide marginal did not need the same care a per-segment computation does: an explicit hairpin-cap guard (S2.3 below), and the multiloop branch energy in the Multi2-side production weight, both verified against the released implementation.

#### S2.1 States and grammar

The algorithm parses each sequence under the stochastic context-free grammar in Table S2, following the Rfold-style decomposition used by CapR and LinearCapR. Main-text Algorithm 1's state types $t$ are the seven non-terminals below; for example, its leaf production is the hairpin closure `StemEnd`$\rightarrow s_n$, its one-non-terminal productions include the internal/bulge closure `StemEnd`$\rightarrow$ $s_m \cdot$`Stem`$\cdot s_n$, and its two-non-terminal production is the multiloop bifurcation `MultiBif`$\rightarrow$ `Multi1`$\cdot$ `Multi2`.

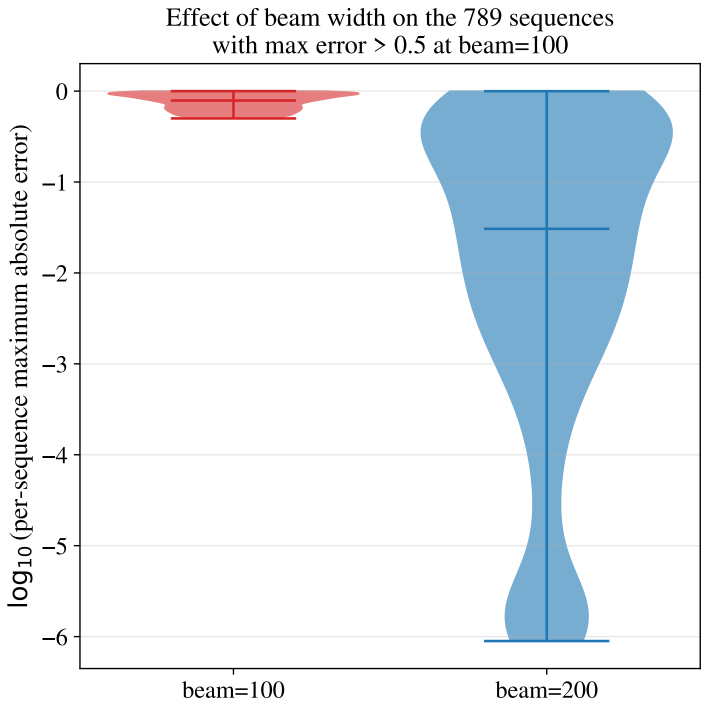


Figure S2: Effect of increasing beam width from 100 to 200 on the 789 sequences with a window error above 0.5 at beam=100: distribution of each sequence's *maximum* absolute error across all requested window lengths ($\log_{10}$ scale; one point per sequence). The median drops from 0.788 (beam=100) to 0.031 (beam=200), but a tail of sequences with maximum error near 1 remains at beam=200.

The production rules are:

$$
\begin{aligned}
\text{Outer} &\rightarrow \epsilon \mid \text{Outer} \cdot a \mid \text{Outer} \cdot \text{Stem} \\
\text{Stem} &\rightarrow b_< \cdot \text{Stem} \cdot b_> \mid b_< \cdot \text{StemEnd} \cdot b_> \\
\text{StemEnd} &\rightarrow s_n \mid s_m \cdot \text{Stem} \cdot s_n \ (m+n>0) \mid \text{Multi} \\
\text{Multi} &\rightarrow s_n \cdot \text{MultiBif} \\
\text{MultiBif} &\rightarrow \text{Multi1} \cdot \text{Multi2} \\
\text{Multi1} &\rightarrow \text{MultiBif} \mid \text{Multi2} \\
\text{Multi2} &\rightarrow \text{Stem} \cdot s_n
\end{aligned}
$$

where $\epsilon$ is the empty string, $a$ is a single unpaired nucleotide, $b_<, b_>$ denote the left/right sides of a base pair, $s_n$ is a run of $n$ consecutive unpaired nucleotides, and "·" denotes concatenation. $t(A \rightarrow B)$ denotes the Boltzmann weight $\exp(-\Delta G/RT)$ of production $A \rightarrow B$. As in the main text (§2.2), hairpin unpaired-run lengths are bounded by $n \leq c_{\text{hairpin}}$, multiloop unpaired-run lengths by $n \leq c_{\text{multi}}$, and internal/bulge closures (the two flanking unpaired runs $m, n$ in StemEnd $\rightarrow s_m \cdot \text{Stem} \cdot s_n$) by the separate, fixed cap $m + n \leq c_{\text{internal}}$ (the ViennaRNA/Turner `MAXLOOP` convention, $c_{\text{internal}} = 30$, not user-configurable); by default $c_{\text{hairpin}} = c_{\text{multi}} = b$, the beam width (the sequence length at beam=0). $\alpha_s(i,j)$ and $\beta_s(i,j)$ denote the inside and outside scores of state $s$ spanning $[i,j]$.

Table S2: Non-terminal symbols in the grammar (identical to LinearCapR's Table 1).

| Name | Description |
|---|---|
| `Outer` | Exterior unpaired region |
| `Stem` | Helix extension |
| `StemEnd` | Terminates a helix into a hairpin, bulge/internal loop, or multiloop region |
| `Multi` | Completed multiloop |
| `MultiBif` | Bifurcation into two or more branches |
| `Multi1` | One-or-more list of branches |
| `Multi2` | One branch followed by a 3′-side run of unpaired bases |

### S2.2 Inside and outside recurrences

The inside scores $\alpha_s(i,j)$ and outside scores $\beta_s(i,j)$ for each state $s$ in Table S2 are computed by the beam-pruned inside–outside recurrence over the grammar above. This recurrence is unchanged from LinearCapR (Otagaki et al., 2026): the same states, the same productions, and the same $b$-best beam pruning after each state's table is filled at a given $j$. We do not reproduce its full pseudocode here, to avoid duplicating a previously published algorithm; readers who want the complete state-by-state recursion should consult LinearCapR's own Methods and Supplementary Material (Otagaki et al., 2026). S2.3 below uses $\alpha_s$ and $\beta_s$ only as already-computed inputs.

### S2.3 Segment-event probabilities

Let $Z = \alpha_{\text{Outer}}(N)$. $B_{\text{Hairpin}}$, $B_{\text{Loop}}$, and $B_{\text{MultiUnpaired}}$ denote the Boltzmann-weighted contributions of hairpin closures, bulge/internal loop closures, and multiloop unpaired segments respectively. We write $p^c_{[s,e]}$ (main text §2.2) as $p([s,e],c)$ to match LinearCapR's own notation. $\alpha_{\text{StemEnd}}(i,j)$ is populated by all three of StemEnd's productions: the hairpin closure $s_n$ (span $[i,j]$, length $j-i+1 \leq c_{\text{hairpin}}$), the internal/bulge closure $s_m \cdot \text{Stem} \cdot s_n$ (unbounded span, capped only via $m+n \leq c_{\text{internal}}$ on the two flanking unpaired runs), and Multi (unbounded span). A retained $[i,j] \in \alpha_{\text{StemEnd}}$ therefore need not itself satisfy the hairpin cap – $j-i+1$ can exceed $c_{\text{hairpin}}$ whenever $\alpha_{\text{StemEnd}}(i,j)$'s dominant contribution came from an internal/bulge or multiloop closure instead – so the hairpin-closure line below includes an explicit guard.

Each event above is a segment $[s,e]$ with probability $p([s,e],c)$; main-text Algorithm 2 adds it, in $O(1)$, to the difference array $D_w$ of every requested window length $w \leq e-s+1$, and one prefix sum per $w$ recovers the window accessibilities. This bounds the cost of each update independently of the segment length; the cost of enumerating segment events is separate.

The two multiloop events above use different weights because they combine different pairs of scores. $\beta_{\text{Multi}}(i,j)$ (used with $\alpha_{\text{MultiBif}}(p,j)$) is already the outside score of a state whose own production, Multi $\rightarrow s_n \cdot$ MultiBif, is the unpaired run itself, so only the unpaired-run weight $B_{\text{MultiUnpaired}}(i,p-1)$ remains to be supplied. $\beta_{\text{Multi2}}(i,j)$ (used with $\alpha_{\text{Stem}}(i,q)$), by contrast, is the outside score of the Multi2 state itself, one level above the production Multi2 $\rightarrow \text{Stem} \cdot s_n$ that connects it to $\alpha_{\text{Stem}}(i,q)$; that production's own weight $t(\text{Multi2} \rightarrow \text{Stem} \cdot s_{j-q})$ – which includes both the per-branch multiloop energy and the unpaired-run energy for $[q+1,j]$ – must therefore be supplied explicitly, rather than the unpaired-run weight alone.

Unlike the other four contexts, the exterior context has no enclosing structure to bound a segment, so it is not obtained from the difference array above. Instead, main text §2.2 computes it

**Algorithm 1** Segment-event probabilities (adapted from LinearCapR, renamed to this paper's $p^c_{[s,e]}$; see text above for the two corrections relative to LinearCapR's own published equations)

**for** each $[i, j] \in \alpha_{\text{StemEnd}}$ **do**
  **if** $j - i + 1 \leq c_{\text{hairpin}}$ **then**
    $p([i, j], H) \mathrel{+}= \frac{1}{Z}\, \beta_{\text{StemEnd}}(i, j) \cdot B_{\text{Hairpin}}(i - 1, j + 1)$
  **end if**
  **for** $p = i + 1$ to $\min(i + c_{\text{internal}}, j - 1)$ **do**
    $p([i, p - 1], B) \mathrel{+}= \frac{1}{Z}\, \alpha_{\text{Stem}}(p, j) \cdot \beta_{\text{StemEnd}}(i, j) \cdot B_{\text{Loop}}(i - 1, j + 1, p, j)$
  **end for**
  **for** $q = \max(j - c_{\text{internal}}, i + 1)$ to $j - 1$ **do**
    $p([q + 1, j], B) \mathrel{+}= \frac{1}{Z}\, \alpha_{\text{Stem}}(i, q) \cdot \beta_{\text{StemEnd}}(i, j) \cdot B_{\text{Loop}}(i - 1, j + 1, i, q)$
  **end for**
  **for** $p = i + 1$ to $\min(i + c_{\text{internal}}, j - 1)$ **do**
    **for** $q = \max(p - i + j - c_{\text{internal}},\ p + 1)$ to $j - 1$ **do**
      $p([i, p - 1], I) \mathrel{+}= \frac{1}{Z}\, \alpha_{\text{Stem}}(p, q) \cdot \beta_{\text{StemEnd}}(i, j) \cdot B_{\text{Loop}}(i - 1, j + 1, p, q)$
      $p([q + 1, j], I) \mathrel{+}= \frac{1}{Z}\, \alpha_{\text{Stem}}(p, q) \cdot \beta_{\text{StemEnd}}(i, j) \cdot B_{\text{Loop}}(i - 1, j + 1, p, q)$
    **end for**
  **end for**
**end for**
**for** each $[p, j] \in \alpha_{\text{MultiBif}}$ **do**
  **for** $i = \max(1, p - c_{\text{multi}})$ to $p - 1$ **do**
    $p([i, p - 1], M) \mathrel{+}= \frac{1}{Z}\, \alpha_{\text{MultiBif}}(p, j) \cdot \beta_{\text{Multi}}(i, j) \cdot B_{\text{MultiUnpaired}}(i, p - 1)$
  **end for**
**end for**
**for** each $[i, q] \in \alpha_{\text{Stem}}$ **do**
  **for** $j = q + 1$ to $\min(N, q + c_{\text{multi}})$ **do**
    $p([q + 1, j], M) \mathrel{+}= \frac{1}{Z}\, \alpha_{\text{Stem}}(i, q) \cdot \beta_{\text{Multi2}}(i, j) \cdot t(\text{Multi2} \to \text{Stem} \cdot s_{j-q})$
  **end for**
**end for**
$p(i, E) = \frac{1}{Z}\, \alpha_{\text{Outer}}(i - 1) \cdot \beta_{\text{Outer}}(i + 1)$ (exterior, per nucleotide)

directly at each requested window $[i, i + w - 1]$ from the boundary partition functions:

$$p([i, i + w - 1], E) = \frac{1}{Z}\, \alpha_{\text{Outer}}(i - 1) \cdot \beta_{\text{Outer}}(i + w) \cdot \exp\Big( - \sum_{k=i}^{i+w-1} E_{\text{ext}}(k)/RT \Big),$$

where $E_{\text{ext}}(k)$ is the per-nucleotide external-loop unpaired energy term, generalizing the per-nucleotide formula above ($w = 1$: $i + w = i + 1$); when this term is 0, the exponential factor is 1 and can be dropped, leaving $\frac{1}{Z}\, \alpha_{\text{Outer}}(i - 1) \cdot \beta_{\text{Outer}}(i + w)$. The paired probability $p(i, S)$ is the complement of the five unpaired-context probabilities, not a member of $\mathcal{C}$. At each position,

$$p(i, S) + \sum_{c \in \mathcal{C}} p(i, c) = 1.$$

### S2.4 What is new in this paper

§S2.2's inside–outside recurrence is unchanged from LinearCapR; §S2.3's segment-event extraction adapts LinearCapR's own published equations, with the two corrections noted above (S2 intro).

LinearCapR's own aggregation stops here: it sums the segment events into per-nucleotide marginals, $\sum_{[s,e]\ni k} p^c_{[s,e]}$, and reports those five numbers per position. This paper's contribution begins at that point: main-text Algorithm 2 takes the same segment events $p^c_{[s,e]}$ derived above and, instead of summing them into a per-nucleotide marginal, adds each event's probability once to a difference array over the range of window-start positions it covers, recovering $P_u(i, i+w-1)$ for every window of every requested length $w$ with one prefix sum per length – the aggregation described in main text §2.2 and Algorithm 2.

### S3. Beam-width sweep on 300–1,000 nt sequences

To check whether increasing the beam width continues to reduce error rather than plateauing, we ran LinearRaccess at beam $\in \{50, 100, 200, 300, 500\}$ (loop caps following the beam) on the 1,770 bpRNA sequences in the 300–1,000 nt range (the three longer length bins of Table 1), against the same Raccess Exact reference. Figure S3 shows the median absolute and opening-energy errors at each beam width, overall and per length bin, pooling all (position, window-length) cells for $w \in \{1, 3, 7, 10, 35\}$. Both decrease monotonically at every tested beam width and in every length bin: overall, the median opening-energy error is 0.71 (beam=50), 0.20 (100), 0.0071 (200), $2 \times 10^{-5}$ (300), and $3 \times 10^{-6}$ kcal/mol (500). Convergence is slower for longer sequences: in the 700–1,000 nt bin the median is still 0.14 kcal/mol at beam=200 and 0.03 kcal/mol at beam=300. The pooled median can also mask tail behavior: the number of sequences with at least one window of absolute error above 0.5 falls from 1,379 (beam=50) to 665 (100), 150 (200), 34 (300), and 3 (500).

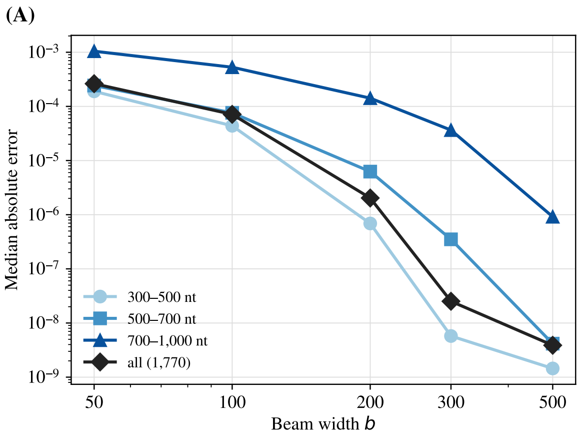


Figure S3: Median absolute error (**A**) and median opening-energy error (**B**) of LinearRaccess against Raccess Exact as a function of beam width, on 1,770 bpRNA sequences (300–1,000 nt), overall and by length bin. Each point pools all (position, window-length) cells, $w \in \{1, 3, 7, 10, 35\}$, across the sequences in that bin.

### S4. Long windows with loop caps tied to the beam

A segment can contain a window $[i, i + w - 1]$ only if its length is at least $w$ and the state defining it survives beam pruning. With the caps tied to the beam, the first condition is met by hairpin and multiloop segments for any $w \le b$ (bulge and internal-loop runs remain bounded by the 30-nt internal-loop limit); with a fixed 30-nt hairpin and multiloop cap, as in earlier versions of LinearRaccess, a 35-nt window could lie only in the exterior loop. The second condition remains: a long unpaired run has fewer base pairs than competing alternatives at the same position, so it has low Boltzmann weight and is disproportionately pruned as $w$ grows. Table S3 shows, on the 1,770-sequence set of S3, that the opening-energy error grows with $w$ at every beam width, and that 35-nt windows need a wider beam than short ones for the same accuracy: at beam=200 the median error is 0.001 kcal/mol at $w$=1 but 0.05 kcal/mol at $w$=35, comparable to that of 1-nt windows at beam=100 (0.035 kcal/mol).

Table S3: Median and 95th-percentile opening-energy error (kcal/mol) of LinearRaccess against Raccess Exact by window length and beam width, on the 1,770 bpRNA sequences of 300–1,000 nt.

| | $w$=1 | 3 | 7 | 10 | 35 |
|---|---|---|---|---|---|
| beam=100, median | 0.035 | 0.097 | 0.22 | 0.32 | 0.92 |
| beam=200, median | 0.001 | 0.003 | 0.008 | 0.012 | 0.052 |
| beam=300, median | <0.001 | <0.001 | <0.001 | <0.001 | <0.001 |
| beam=500, median | <0.001 | <0.001 | <0.001 | <0.001 | <0.001 |
| beam=100, p95 | 1.33 | 1.92 | 2.69 | 3.14 | 5.57 |
| beam=200, p95 | 0.39 | 0.64 | 1.02 | 1.24 | 2.61 |
| beam=300, p95 | 0.071 | 0.13 | 0.24 | 0.32 | 0.95 |
| beam=500, p95 | 0.002 | 0.004 | 0.008 | 0.010 | 0.037 |

### S5. Comparison with LinearSampling

We compared LinearRaccess with LinearSampling (Zhang et al., 2023) under the Turner 2004 model on two sequence sets: 52 bpRNA sequences (13 per Table 1 length bin, of which the 26 longer than 500 nt are reported below, since for sequences no longer than the beam width LinearRaccess prunes nothing), and 20 bpRNA sequences of 1,000–3,541 nt (5 per bin of 1,000–1,500, 1,500–2,000, 2,000–3,000, and 3,000–4,381 nt, in file order). Both tools were run at beam widths 100–500, LinearSampling with $k = 10^6$ samples, and scored against LinearRaccess in exact mode (beam=0, exact log-sum-exp). This reference agrees with Raccess Exact under Turner 1999 (main text §3.1) and, under Turner 2004, with LinearSampling's own exact sampling: on three sequences of 288–654 nt, LinearSampling with beam 0 and $2 \times 10^5$ samples matched it within sampling noise at all 9,555 windows (totals and each loop context) with accessibility $\geq 10^{-3}$, the largest deviation being 4.1 standard errors. Runtime and peak memory were measured one process at a time on a single core; LinearSampling's time includes counting windows in its samples, since that is what produces accessibility.

Table S4 gives the median relative error by the exact accessibility's decade. On the 524–973 nt sequences LinearRaccess at beam=500 is more accurate than LinearSampling at every decade, at similar time. On the 1,000–3,541 nt sequences, LinearSampling at the same beam width is more accurate for windows with accessibility above $10^{-4}$, whereas LinearRaccess is more accurate below it, where LinearSampling's error approaches 100% as windows are sampled rarely or never. Since both methods evaluate the ensemble that survives beam pruning, the difference at high accessibility reflects their pruning: LinearRaccess's grammar splits multiloops over more state types, each pruned to $b$ states. Table S5 gives the corresponding opening-energy view: LinearSampling assigns an infinite opening energy (accessibility exactly zero) to a growing fraction of windows as $w$ grows – about 20% at $w$=10 and over 95% at $w$=35 – which leaves its within-sequence ranking of 35-nt windows almost uninformative, whereas LinearRaccess returned a finite opening energy for every window of both sets.

Table S4: Median relative error $|P_{\text{approx}} - P_{\text{exact}}|/P_{\text{exact}}$ by decade of the exact accessibility (all windows, $w \in \{1, 3, 7, 10, 35\}$), overall median over windows with $P_{\text{exact}} \geq 10^{-8}$, number of sequences with some window of absolute error above 0.5, and single-core total time and peak memory. LinearSampling uses $k = 10^6$.

| | Median relative error by $P_{\text{exact}}$ | | | | | | | | | |
|---|---|---|---|---|---|---|---|---|---|---|
| Method | $\geq 10^{-1}$ | $[10^{-2}, 10^{-1})$ | $[10^{-3}, 10^{-2})$ | $[10^{-4}, 10^{-3})$ | $[10^{-6}, 10^{-4})$ | $[10^{-8}, 10^{-6})$ | Overall | Fail | Time (s) | Memory (MB) |
| *26 sequences, 524–973 nt* | | | | | | | | | | |
| LinearRaccess $b$=200 | 0.024 | 0.11 | 0.14 | 0.17 | 0.21 | 0.26 | 0.12 | 6 | 129 | 93 |
| LinearRaccess $b$=500 | 0.0003 | 0.0005 | 0.0004 | 0.0006 | 0.0007 | 0.0011 | 0.0004 | 0 | 279 | 153 |
| LinearSampling $b$=100 | 0.053 | 0.23 | 0.30 | 0.40 | 0.70 | 1.00 | 0.34 | 11 | 238 | 48 |
| LinearSampling $b$=500 | 0.0007 | 0.006 | 0.018 | 0.061 | 0.24 | 1.00 | 0.017 | 0 | 284 | 120 |
| *20 sequences, 1,000–3,541 nt* | | | | | | | | | | |
| LinearRaccess $b$=200 | 0.17 | 0.56 | 0.60 | 0.66 | 0.71 | 0.81 | 0.52 | 19 | 471 | 444 |
| LinearRaccess $b$=500 | 0.028 | 0.13 | 0.14 | 0.16 | 0.20 | 0.23 | 0.11 | 8 | 1,993 | 909 |
| LinearSampling $b$=200 | 0.085 | 0.35 | 0.41 | 0.50 | 0.85 | 1.00 | 0.45 | 17 | 884 | 255 |
| LinearSampling $b$=500 | 0.008 | 0.038 | 0.062 | 0.14 | 0.45 | 1.00 | 0.098 | 2 | 1,150 | 468 |

### S6. Comparison with DeepRaccess

The published DeepRaccess model predicts the opening energy of 35-nt windows only; its teacher is Raccess Window ($W = 100$) run with CONTRAfold parameters instead of Turner energies (Hara et al., 2023). We reproduced this teacher exactly: the example teacher values distributed

Table S5: Opening-energy view of the same comparison: median opening-energy error (kcal/mol, over windows where both values are finite), percentage of windows with infinite opening energy (approximate accessibility exactly zero, exact accessibility positive), and median within-sequence Spearman correlation with the exact opening energies.

| | $w$=1 | | | $w$=10 | | | $w$=35 | | |
|---|---|---|---|---|---|---|---|---|---|
| Method | error | ∞ (%) | Spearman | error | ∞ (%) | Spearman | error | ∞ (%) | Spearman |
| *26 sequences, 524–973 nt* | | | | | | | | | |
| LinearRaccess $b$=200 | 0.027 | 0 | 0.991 | 0.13 | 0 | 0.964 | 0.34 | 0 | 0.945 |
| LinearRaccess $b$=500 | <0.001 | 0 | 1.000 | <0.001 | 0 | 1.000 | 0.001 | 0 | 0.999 |
| LinearSampling $b$=200 | 0.014 | 0.2 | 0.998 | 0.12 | 20.4 | 0.966 | 0.71 | 99.7 | 0.12 |
| LinearSampling $b$=500 | 0.001 | 0.1 | 1.000 | 0.049 | 18.4 | 0.984 | 0.76 | 99.6 | 0.14 |
| *20 sequences, 1,000–3,541 nt* | | | | | | | | | |
| LinearRaccess $b$=200 | 0.14 | 0 | 0.872 | 0.61 | 0 | 0.840 | 1.32 | 0 | 0.867 |
| LinearRaccess $b$=500 | 0.024 | 0 | 0.982 | 0.13 | 0 | 0.968 | 0.30 | 0 | 0.961 |
| LinearSampling $b$=200 | 0.087 | 0.4 | 0.928 | 0.38 | 26.4 | 0.868 | 0.54 | 95.6 | 0.17 |
| LinearSampling $b$=500 | 0.013 | 0.3 | 0.998 | 0.10 | 21.0 | 0.979 | 0.32 | 95.2 | 0.16 |

with DeepRaccess match `run_raccess_contrafold -access_len=35 -max_span=100` to all printed digits. Because CONTRAfold scores and Turner 1999 energies are on different scales, we scored each method against the model it approximates, at $w = 35$ on the 12,872 sequences of Table 1 (Table S6). DeepRaccess's median error against its teacher is 0.4–0.6 kcal/mol in every length bin, and its within-sequence ranking agreement varies widely between sequences (median Spearman 0.88, 5th percentile 0.45); LinearRaccess's error against Raccess Exact is far smaller up to 700 nt and grows with length to 0.62 kcal/mol at 700–1,000 nt, with median Spearman 1.00 (5th percentile 0.98). Because the two methods use different underlying energy models, these error values are not directly comparable as measures of physical accessibility accuracy. DeepRaccess's predictions are also local, on two counts: its teacher restricts base pairs to a span of $W = 100$, and its preprocessing splits a sequence longer than 440 nt into 440-nt segments overlapping by 110 nt, predicts each segment separately, and joins them after discarding 55 nt at each overlapping end; LinearRaccess processes each sequence whole and without a span limit. DeepRaccess is much faster: over the 52 sequences of S5, 6.1 s in total on one CPU thread (6.7 s on one NVIDIA H100 GPU, one sequence at a time), versus 160 s for LinearRaccess (beam=200); in batches on the GPU, all 12,872 sequences took 31 s.

Table S6: Median opening-energy error (kcal/mol) at $w = 35$ on the 12,872 sequences of Table 1, each method against the model it approximates; for reference, the teacher itself against CONTRAfold without a span limit.

| Comparison | 100–300 | 300–500 | 500–700 | 700–1,000 | All |
|---|---|---|---|---|---|
| DeepRaccess vs. its teacher (CONTRAfold, $W$=100) | 0.59 | 0.48 | 0.44 | 0.39 | 0.53 |
| LinearRaccess ($b$=200) vs. Raccess Exact (Turner 1999) | <0.001 | 0.019 | 0.19 | 0.62 | <0.001 |
| Teacher vs. CONTRAfold without span limit | 0.15 | 0.97 | 0.91 | 0.80 | 0.38 |

### S7. Accessibility by loop context

LinearRaccess reports each window's accessibility split into the five loop contexts, $P_u^c(i, w)$ (main text Eq. 5), with no change in cost (option `-byloop`; Table S7, last column). We compared these values with LinearSampling's, obtained by assigning each unpaired run of every sampled structure the context of its innermost enclosing base pair, on the sequence sets of S5 against LinearRaccess in exact mode (Table S7). Splitting a sampling estimate by context divides its counts, so windows with small per-context accessibility are often sampled zero times (11–66% of windows, depending on context), whereas LinearRaccess returns a nonzero value for every window and, at the same beam width, a smaller error in most contexts.

Table S7: Accuracy of per-context window accessibility $P_u^c$ against LinearRaccess in exact mode, over windows with exact $P_u^c \geq 10^{-8}$ ($w \in \{1, 3, 7, 10, 35\}$). Each cell gives the median relative error, followed in parentheses by the percentage of those windows estimated as exactly zero; the last column is the single-core total time. LinearSampling uses $k = 10^6$ samples.

| Method | Hairpin | Bulge | Internal | Multiloop | Exterior | Time (s) |
|---|---|---|---|---|---|---|
| *26 sequences, 524–973 nt* | | | | | | |
| LinearRaccess $b$=200 | 0.18 (0) | 0.19 (0) | 0.16 (0) | 0.27 (0) | 0.48 (0) | 131 |
| LinearRaccess $b$=500 | 0.001 (0) | 0.001 (0) | 0.001 (0) | 0.001 (0) | 0.001 (0) | 283 |
| LinearSampling $b$=200 | 0.25 (17) | 0.45 (30) | 0.28 (23) | 0.21 (12) | 1.00 (47) | 295 |
| LinearSampling $b$=500 | 0.07 (16) | 0.19 (28) | 0.09 (21) | 0.04 (11) | 0.87 (45) | 306 |
| *20 sequences, 1,000–3,541 nt* | | | | | | |
| LinearRaccess $b$=200 | 0.71 (0) | 0.69 (0) | 0.63 (0) | 0.81 (0) | 1.00 (0) | 461 |
| LinearRaccess $b$=500 | 0.17 (0) | 0.18 (0) | 0.14 (0) | 0.23 (0) | 0.46 (0) | 1,940 |
| LinearSampling $b$=200 | 0.75 (22) | 0.91 (32) | 0.76 (24) | 0.74 (17) | 1.00 (66) | 904 |
| LinearSampling $b$=500 | 0.21 (17) | 0.40 (27) | 0.24 (20) | 0.18 (13) | 1.00 (54) | 1,165 |
